# Fifteen shades of clay: distinct microbial community profiles obtained from bentonite samples by cultivation and direct nucleic acid extraction

**DOI:** 10.1101/2021.07.13.452221

**Authors:** Melody A. Vachon, Katja Engel, Rachel C. Beaver, Greg W. Slater, W. Jeffrey Binns, Josh D. Neufeld

## Abstract

Characterizing the microbiology of swelling bentonite clays can help predict the long-term behaviour of deep geological repositories (DGRs), which are proposed as a solution for the management of used nuclear fuel worldwide. Such swelling clays represent an important component of several proposed engineered barrier system designs and, although cultivation-based assessments of bentonite clay are routinely conducted, direct nucleic acid detection from these materials has been difficult due to technical challenges. In this study, we generated direct comparisons of microbial abundance and diversity captured by cultivation and direct nucleic acid analyses using 15 reference bentonite clay samples. Regardless of clay starting material, the corresponding profiles from cultivation-based approaches were consistently associated with phylogenetically similar sulfate-reducing bacteria, denitrifiers, aerobic heterotrophs, and fermenters, demonstrating that any DGR-associated growth may be consistent, regardless of the specific bentonite clay starting material selected for its construction. Furthermore, dominant nucleic acid sequences in the as-received clay microbial profiles did not correspond with the bacteria that were enriched or isolated in culture. Few core taxa were shared among cultivation and direct nucleic acid analysis profiles, yet those in common were primarily affiliated with *Streptomyces, Micrococcaceae, Bacillus*, and *Desulfosporosinus* genera. These putative desiccation-resistant bacteria associated with diverse bentonite clay samples can serve as targets for experiments that evaluate microbial viability and growth within DGR-relevant conditions. Our data will be important for global nuclear waste management organizations, demonstrating that identifying appropriate design conditions with suitable clay swelling properties will prevent growth of the same subset of clay-associated bacteria, regardless of clay origin or processing conditions.

## Introduction

Numerous countries have amassed used nuclear fuel resulting from power generation over several decades. Although this spent fuel is held within temporary above-ground storage, it will remain radioactive for millennia and requires a permanent solution (1). In accordance with international consensus on best-practices, many countries are in various stages of planning, designing, and constructing a deep geological repository (DGR) for sustainable isolation and management of used nuclear fuel (2,3). Even though design features vary from country to country, all DGR design plans propose to bury used fuel within a stable geological formation, surrounded by multiple engineered barriers (e.g., Figure S1). In Canada, Sweden, and Finland, for example, engineered barriers include storing used nuclear fuel bundles within carbon steel used fuel containers, for strength, with either an integrally bonded copper coating (e.g., Canada) or a self-supporting outer copper shell (e.g., Sweden and Finland) for corrosion resistance (4,5). An additional engineered barrier involves surrounding used fuel containers with highly compacted bentonite clay, which swells when saturated. This swelling action serves to decrease water activity and microbial growth, while restricting transport of oxidants toward the used fuel container and migration of radionuclides in the unlikely event of container failure (6). Bentonite clay thus serves as an important engineered barrier system component within the natural barrier of a stable host rock. Together, the combination of natural and engineered barrier components of a DGR are intended to isolate and contain nuclear waste through multiple glaciation cycles until relatively safe levels of radiation are reached within approximately one million years (4).

Because microorganisms are present within mined materials, such as bentonite clay, and are introduced during storage and processing, several studies have evaluated potential microbial impacts on engineered barrier system components related to copper corrosion, biofilm formation, radionuclide transport, transformation of clay minerals, and gas production (6,7). One of the primary microbiological concerns relevant to long-term containment of nuclear waste is microbiologically influenced corrosion. Such corrosion might occur as active microorganisms release metabolites that directly or indirectly cause metal corrosion (8,9). For example, sulfate-reducing bacteria (SRB) generate hydrogen sulfide (H_2_S), which can be corrosive to metals such as copper and steel (8,10–13). Heterotrophic bacteria are also relevant for DGR safety assessments because fermentation-associated hydrogen gas (H_2_) and acetate production may promote sulfate-reduction by SRB through increased electron donor availability (12,14). Production of gases by microorganisms like denitrifiers, methanogens, and methanotrophs is important to consider because the gases could lead to the formation of fissures in compacted clay, which may potentially allow transport of microorganisms, microbially produced compounds, or radionuclides in an improbable escape from used fuel containers (6,15). Overall, evaluating the potential for microbial growth and activity within natural and engineered barrier components is an important priority for predicting DGR stability and identifying conditions that minimize or prevent microbial viability over geological timeframes.

Given that DGR-associated microorganisms will be derived from natural or engineered barrier components, an important predicate to modeling the potential impacts of microorganisms on DGR stability is developing an understanding of the microbiota naturally present in these components. Several studies have used cultivation-based approaches to enumerate SRB within bulk clay samples or clay subjected to experimental treatments under DGR-relevant conditions. In these studies, most probable number (MPN) tubes with sulfate-containing medium are used to estimate SRB abundances (13,16–24). Heterotrophic bacteria from bentonite clay are typically grown on R2A medium (16,18,19,21,22,24–28) because of reduced nutrient concentrations (29), which may better mimic limited nutrient availability expected for clay samples. Detection of both aerobic and anaerobic heterotrophs are relevant for these studies because the DGR is likely to shift from oxic to anoxic conditions after a relatively short period of time (16,18–20,24). Microorganisms capable of respiring nitrate, such as denitrifiers, have also been studied in the context of the DGR because of the potential impacts of nitrogen oxide and dinitrogen gas production (10,21,30).

Despite progress in quantifying and characterizing culturable microorganisms from bentonite clays, the extent to which cultivation represents all viable and relatively abundant clay microorganisms is unknown. Enumeration with traditional cultivation approaches limits detection to only those microorganisms that can grow under specific laboratory conditions. Furthermore, cultivation approaches overlook viable but non-culturable bacteria and fastidious or slow-growing microorganisms. Although community profiling methods, based on extracted nucleic acids, can help assess cultivation bias, sequencing of amplified 16S rRNA genes from bentonite clays has seldom been reported, presumably due to low nucleic acid yields that result from low biomass samples and the sorption of DNA onto charged montmorillonite clay layers (31). Despite these limitations, a protocol for successful extraction of DNA from bentonite clay samples has recently been validated(22), allowing us to investigate whether dominant ASVs detected in as-received bentonite clays were the same as those identified through culture-dependent methods, and whether the taxa we culture differ based on clay composition, origin, and storage conditions. In addition to describing microbial community composition, clay microorganisms were quantified with cultivation, quantitative PCR (qPCR) of 16S rRNA genes, and phospholipid fatty acid (PLFA) analysis. Identifying a “core microbiome” and core culturable community subset from diverse clay samples and production lots represents an important step toward validating the choice of dry bentonite starting material in future experiments that will help identify suitable DGR conditions for preventing microbial activity and growth over geological timeframes.

## Results and Discussion

### Microbial heterogeneity in bentonite clays

Assessing microbial profile heterogeneity within diverse commercially available bentonite clays is critical for predicting the microbial growth and activity that may occur within a DGR. In this study, 15 industrially processed bentonite clay samples were sourced from Canada, Greece, India, and the United States of America, with varying production dates, lot numbers, colours, and proportions of exchangeable cations (Table 1; Figure 1). Although the manufacturing and storage process for each of our samples remain relatively unknown, studying a wide range of industrially mined and processed bentonites is essential to capture all possible variations that may be included in future large-scale DGR design and construction. The as-received clay samples were all relatively dry, with moisture contents ranging from 5-16% and water activities between 0.26-0.70 (Figure S2), all well below the water activity threshold of 0.96 considered suitable for microbial growth in bentonite clay (24,32).

**Table 1.** Diverse samples of bentonite clay from different countries.

| Sample | Clay type | Exchangeable cation | Origin | Manufacturer | Production date<sup>a</sup> | Lot number<sup>a</sup> |
| --- | --- | --- | --- | --- | --- | --- |
| MX1 | MX-80 bentonite | Na<sup>+</sup> | Wyoming, USA | American Colloid Company | Jun 2015 | 65275772 |
| MX2 | MX-80 bentonite | Na<sup>+</sup> | Wyoming, USA | American Colloid Company | Nov 2016 | 116315319 |
| MX3 | MX-80 bentonite | Na<sup>+</sup> | Wyoming, USA | American Colloid Company | Mar 2017 | 37324182 |
| MX4 | MX-80 bentonite | Na<sup>+</sup> | Wyoming, USA | American Colloid Company | Mar 2017 | 37324184 |
| MX5 | MX-80 bentonite | Na<sup>+</sup> | Wyoming, USA | American Colloid Company | Mar 2017 | 37324190 |
| MX6 | MX-80 bentonite | Na<sup>+</sup> | Wyoming, USA | American Colloid Company | NA | NA |
| MX7 | MX-80 bentonite | Na<sup>+</sup> | Wyoming, USA | American Colloid Company | NA | NA |
| MX8 | MX-80 bentonite | Na<sup>+</sup> | Wyoming, USA | American Colloid Company | Jun 2015 | 65275768 |
| MX9 | MX-80 bentonite | Na<sup>+</sup> | Wyoming, USA | American Colloid Company | NA | NA |
| AB1 | Asha bentonite | Na<sup>+</sup> | Kutch, India | Ashapura Minechem Co. | NA | NA |
| CC1 | Sodium bentonite | Na<sup>+</sup> | Saskatchewan, Canada | Canadian Clay Products Inc. | NA | NA |
| CC2 | Sodium bentonite | Na<sup>+</sup> | Saskatchewan, Canada | Canadian Clay Products Inc. | NA | NA |
| DC1 | Deponit Ca-N | Ca<sup>2+</sup> | Milos, Greece | S&B Industrial Minerals SA | NA | NA |
| IR1 | IBECO-RWC | Ca<sup>2+</sup> | Milos, Greece | S&B Industrial Minerals SA | NA | NA |
| NB1 | National standard | Na<sup>+</sup> | Wyoming, USA | Opta Minerals Inc. | NA | NA |

<sup>a</sup>NA, data not available

**Figure 1.**
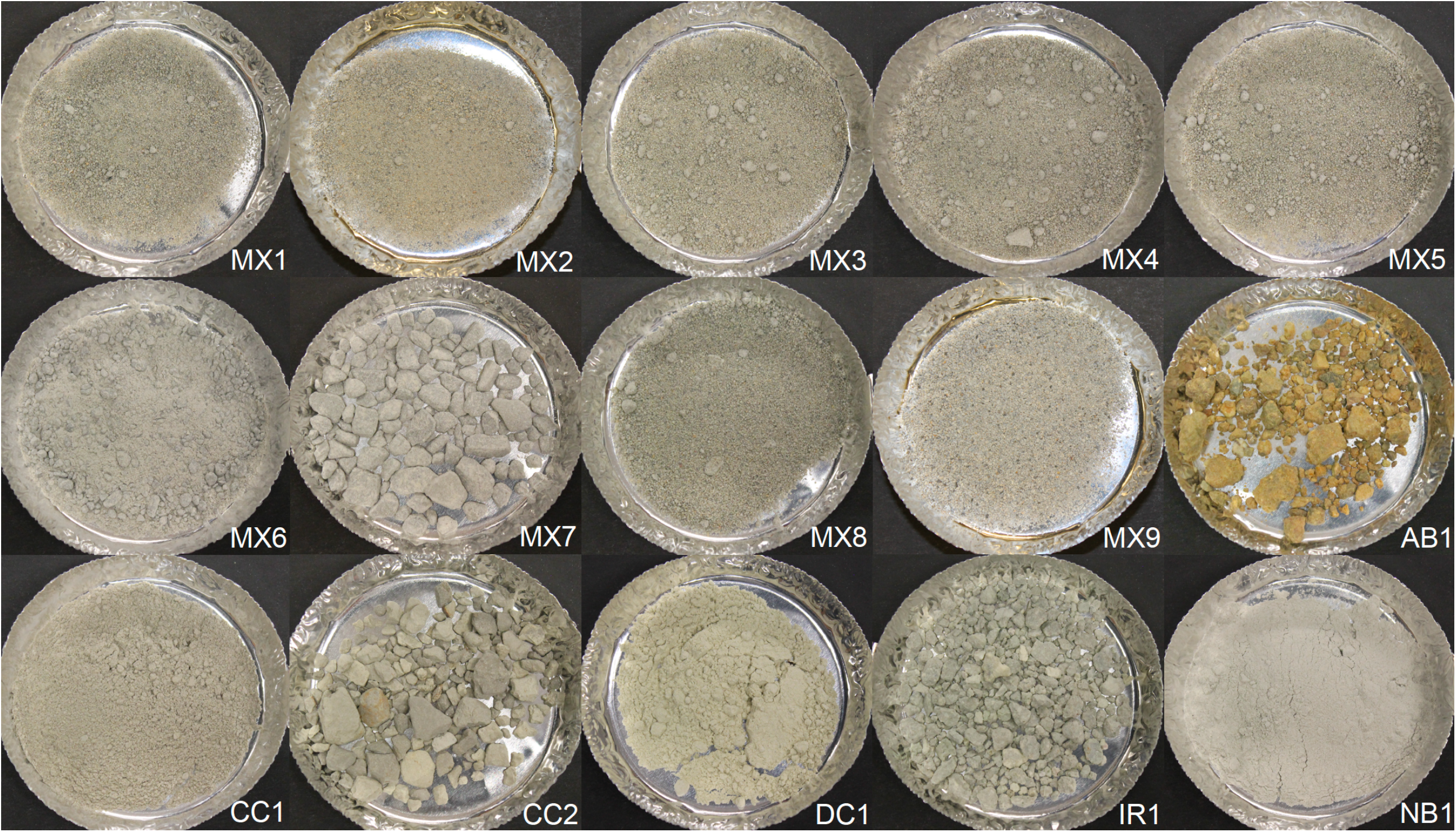
Photographs of the fifteen diverse samples of clay used in this study. Originally coarse samples (AB1, CC2, IR1, and MX7) were ground to smaller grain sizes before use in experiments.

We compared culturable aerobic and anaerobic heterotrophs, SRB, and denitrifying bacteria from all samples. As well, quantification of biomarkers (i.e., 16S rRNA genes and PLFA) provided culture-independent enumerations and taxonomic profiles. Cultivation of microorganisms yielded lower abundance estimates than those obtained by DNA and PLFA quantification (Figure 2), in most cases by orders of magnitude. Higher 16S rRNA gene copy numbers compared to culturable abundance estimations may be due to multiple 16S rRNA gene copies per genome or detection of “relic” DNA within samples. Most culturable aerobic heterotroph abundances ranged from 10^2^ to 10^4^ colony forming units per gram dry weight (CFU/gdw), which is within the range previously reported for bentonite clays (10^2^ to 10^5^ CFU/g; 16, 18, 25, 27). Eleven samples contained average aerobic heterotroph abundances below the limit of plate count quantification (i.e., 2500 CFU/gdw, based on 25 colonies per plate minumum). Anaerobic heterotroph abundance averages were lower than aerobic heterotroph averages for all samples (Figure 2). All anaerobic heterotroph enumerations were below the lower limit of plate count quantification. Average most probable number estimates of SRB from bentonite samples was 63 ± 87 MPN/gdw, comparable to previously studied as-received bentonite samples with SRB abundances of up to 42 MPN/g (18,24,26). The average estimated abundance of culturable denitrifying bacteria was 57 ± 36 MPN/gdw. Overall, sample AB1 had the highest average abundance estimates determined by all enumeration methods, with the exception of PLFA analysis, for which it had the second highest abundance estimate (Figure 2).

**Figure 2.**
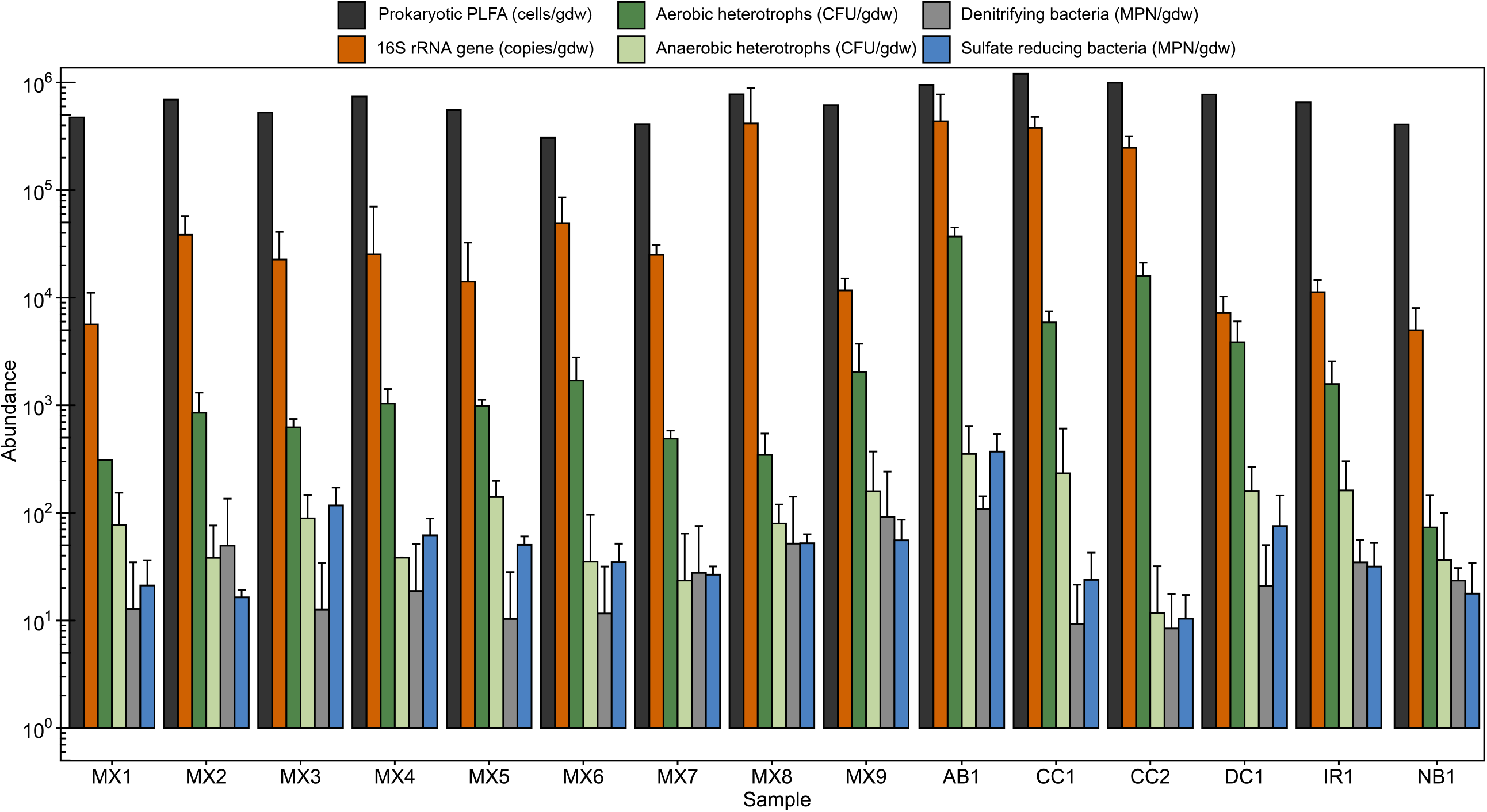
Microbial abundances estimated through culture-dependent and culture-independent methods and normalized to gram dry weight (gdw) using moisture content measurements. Error bars show the standard deviation of triplicate culture-based enumerations and the standard deviation of duplicate qPCR amplifications. Measurements for aerobic and anaerobic heterotrophs below 2.5 × 10^3^ CFU/gdw are below the lower limit of plate count quantification (i.e., 25 colonies per plate) but are shown here for comparison nonetheless. The PLFA analyses were performed in singlicate (duplicate for AB1 and IR1) and only the predicted prokaryotic abundances are reported here for comparison.

Quantification of PLFA from clay samples yielded the highest estimates of microbial cell abundance compared to the other enumeration methods (Figure 2). Analysis of PLFA has been used previously to detect eukaryotic biomarkers in clay (33) and these biomarkers were also detected in the present study, ranging from 10^3^ to 10^4^ cells/gdw, and accounting for 1.6% to 11.0% of the PLFA-associated predicted cell abundances (Figure S3). For all clay samples, the PLFA abundances ranged from 17-64 pmol/gdw, which corresponds to calculated estimates of 10^5^-10^6^ cells/gdw. The cell abundances based on PLFA are comparable to previous reports of PLFA in bentonite and Opalinus Clays that presented quantities of 10^6^ cells/g (24,33,34). These previous experiments also identified that the microbial cell abundance estimates based on PLFA exceeded those based on cultivation by ∼1000-fold (24,33,34). The reasons that PLFA analysis may be associated with relatively high biomass estimates may be related to preservation of PLFA within the clay matrix, as also occurs for DNA. In most environmental samples, PLFA are assumed to degrade within days to weeks of cell death due to biological recycling, thus the remaining PLFA represent viable biomass. However, in clay environments, it has been suggested that PLFA turn-over rates can vary due to environmental conditions like pH (35) and adsorption to clay surfaces (34,36), or by preservation within the clay matrix (33,34). Overall, the quantity of background “noise” that may be detected relative to the abundance of living microorganisms remains unknown.

Microbial community profiles were generated for all clay samples using high-throughput sequencing of 16S rRNA genes. The microbial profiles generated from replicate extractions of the same clay sample usually grouped together within ordination space (Figure 3). The duplicates of samples IR1, DC1, and NB1 exhibited higher dissimilarity compared to other duplicate extraction pairs. Apparent dissimilarity among duplicate clay samples may be due to clay sample heterogeneity or variable extraction efficiencies and detection limits of particular bentonite samples (22,37). Of the microbial community profiles detected in the clay samples, those from MX-80 bentonite clays with the same production date grouped together despite different lot numbers (Figure 3). Previous analysis of bentonite clays also revealed similarities between the microbial community profiles of clays from similar production dates (22). Nonetheless, microbial community profiles of clay samples from March 2017 (MX3, MX4, and MX5) grouped closer together whereas samples from June 2015 (MX1 and MX8) showed more separation. Bentonite clay samples from Wyoming, USA also did not group separately from bentonite from different locations (e.g., Saskatchewan, Kutch, and Milos).

**Figure 3.**
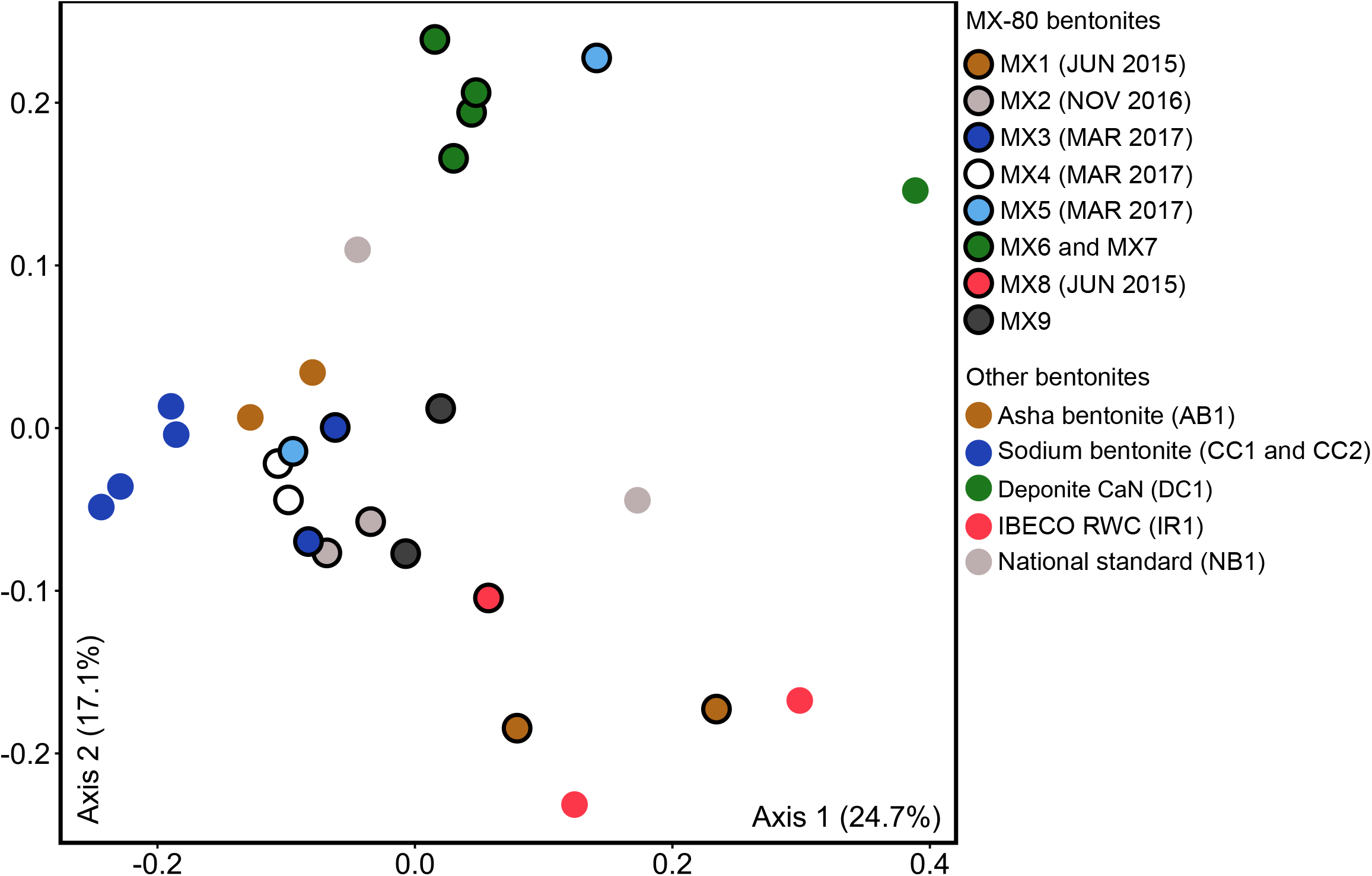
Principle-coordinate analysis (PCoA) ordination based on the weighted UniFrac distance metric of 16S rRNA gene sequences generated by direct DNA extraction from as-received clays. Replicates of DC1 and MX8 were removed during normalization due to low read counts.

Distinct microbial 16S rRNA gene profiles were associated with each tested clay (Figure 4). Many amplicon sequence variants (ASVs) were unique to a single clay sample, but several were more broadly detected across clay samples, such as those affiliated with *Streptomyces, Sphingomonas, Thiobacillus*, and *Xanthomonas*. The potential contaminant ASVs that were flagged by Decontam were detected with low relative abundance withinin samples and were removed from the data (Figure S4). Overall, most ASVs detected in bentonite clay samples were associated with members of the *Actinobacteria* and *Proteobacteria* phyla, similar to profiles previously generated from other MX-80 bentonite clay samples (22,38).

**Figure 4.**
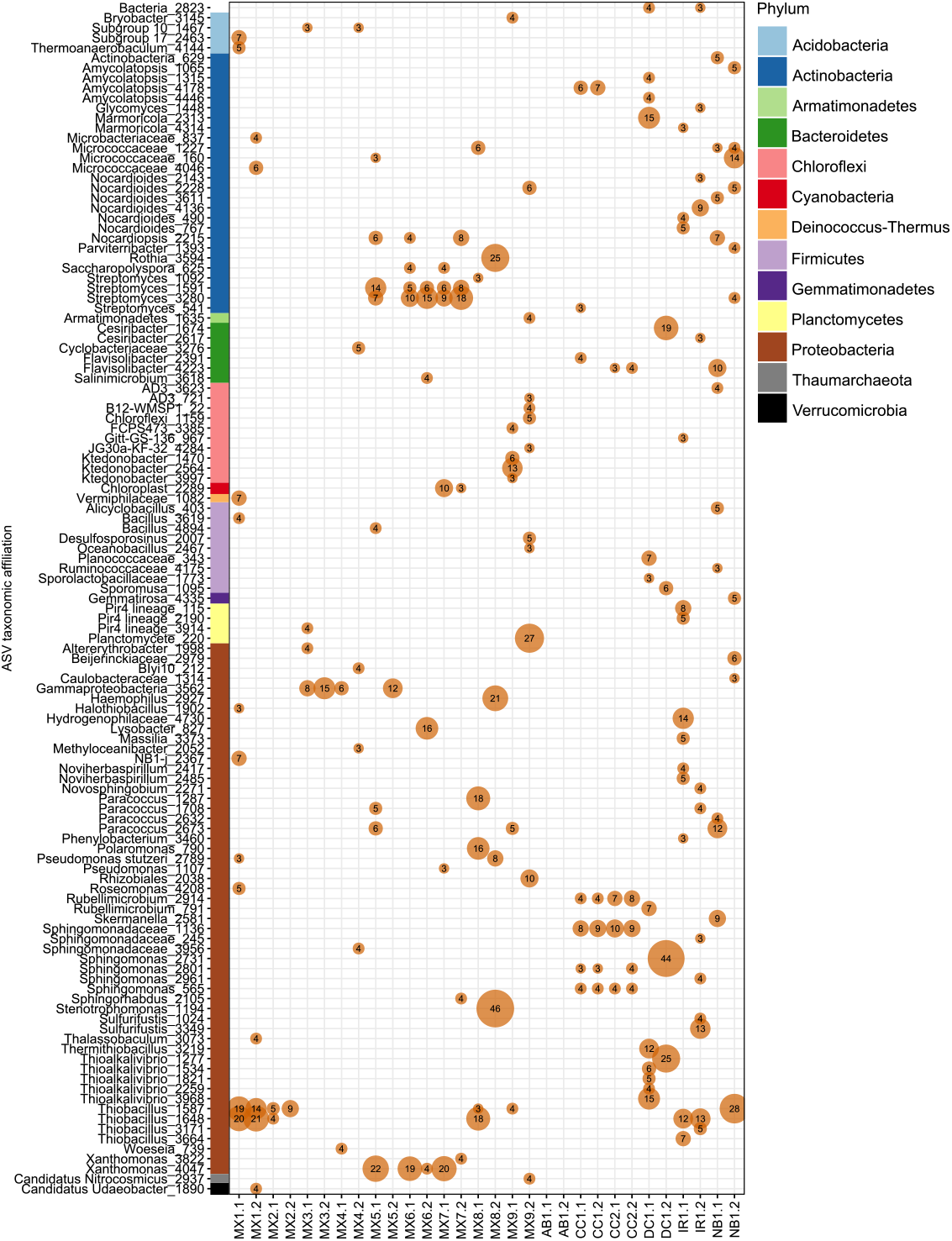
The ASV-level taxa affiliated with as-received clay samples. Phylum of each ASV is indicated with the coloured rectangular bar along the y-axis. Duplicates (denoted as “1” and “2”) from direct DNA extractions of each clay sample are shown, and ASVs listed are ≥ 3% relative abundance in samples. For ASV labels, we report the lowest taxonomic ranks that have confidence values above the default 0.7 threshold.

### Comparison of cultivation and nucleic acid isolation approaches

By generating parallel cultivation-dependent and cultivation-independent microbial profiles, this study is the first to directly compare the microorganisms cultivated from industrially processed bentonite clay to those detected by direct DNA extraction, PCR, and 16S rRNA gene sequencing. Overall, the dominant taxa associated with 16S rRNA gene profiles generated from as-received bentonite clays do not reflect the mircoorganisms that responded to cultivation, and the taxa that were cultivated were very similar, regardless of the starting clay material.

The differences in microbial community profiles generated using cultivation-independent and cultivation-dependent methods are reflected by separate grouping of these two sample types in ordination space (Figure 5), and can be attributed to the culture-based growth of only a subset of the taxa accounted for in the DNA extracted directly from the clay samples (Figure 6). Notably, the absence of many culture ASVs in the dry clay may be due to adsorption of DNA to the clay matrix, incomplete lysing of viable spores during DNA extraction or insufficient sequencing depth. The microorganisms that were detected in cultures were commonly from the orders *Bacillales* and *Clostridiales*. Although similar orders were detected from the same type of enrichment across all samples, cultures from each sample had unique microbial profiles at the ASV level (Figures S5, S6, and S7). Here, and in previous research, several of the most frequently detected ASVs from cultures were affiliated with the *Bacillus* and *Clostridium* genera (13,18). Dominant taxa detected in cultures were low in abundance or even absent in the microbial profiles of the corresponding clay samples (Figure 4).

**Figure 5.**
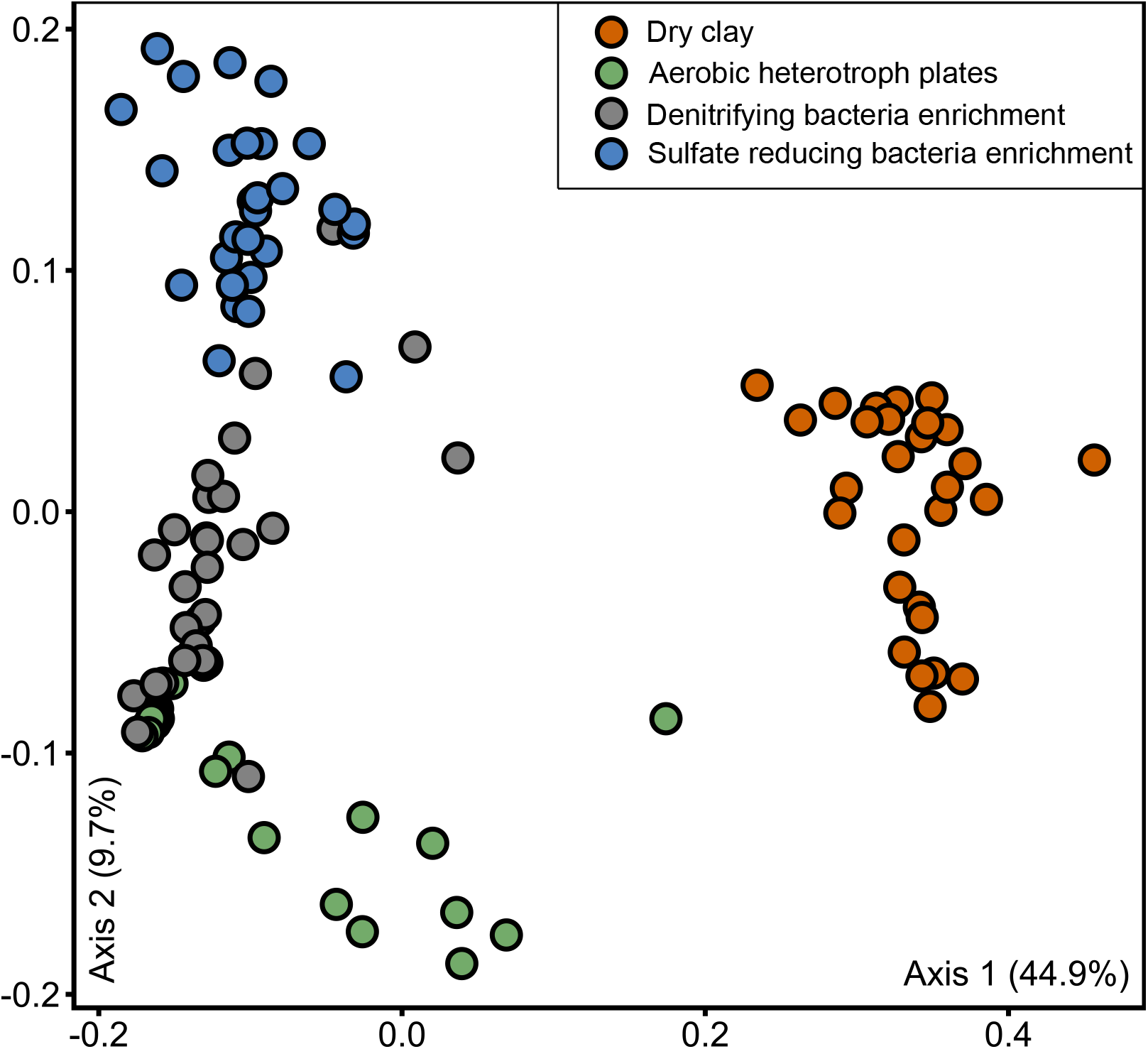
Weighted UniFrac principle-coordinate analysis (PCoA) ordination of 16S rRNA gene sequences generated from as-received clays and enrichment cultures.

**Figure 6.**
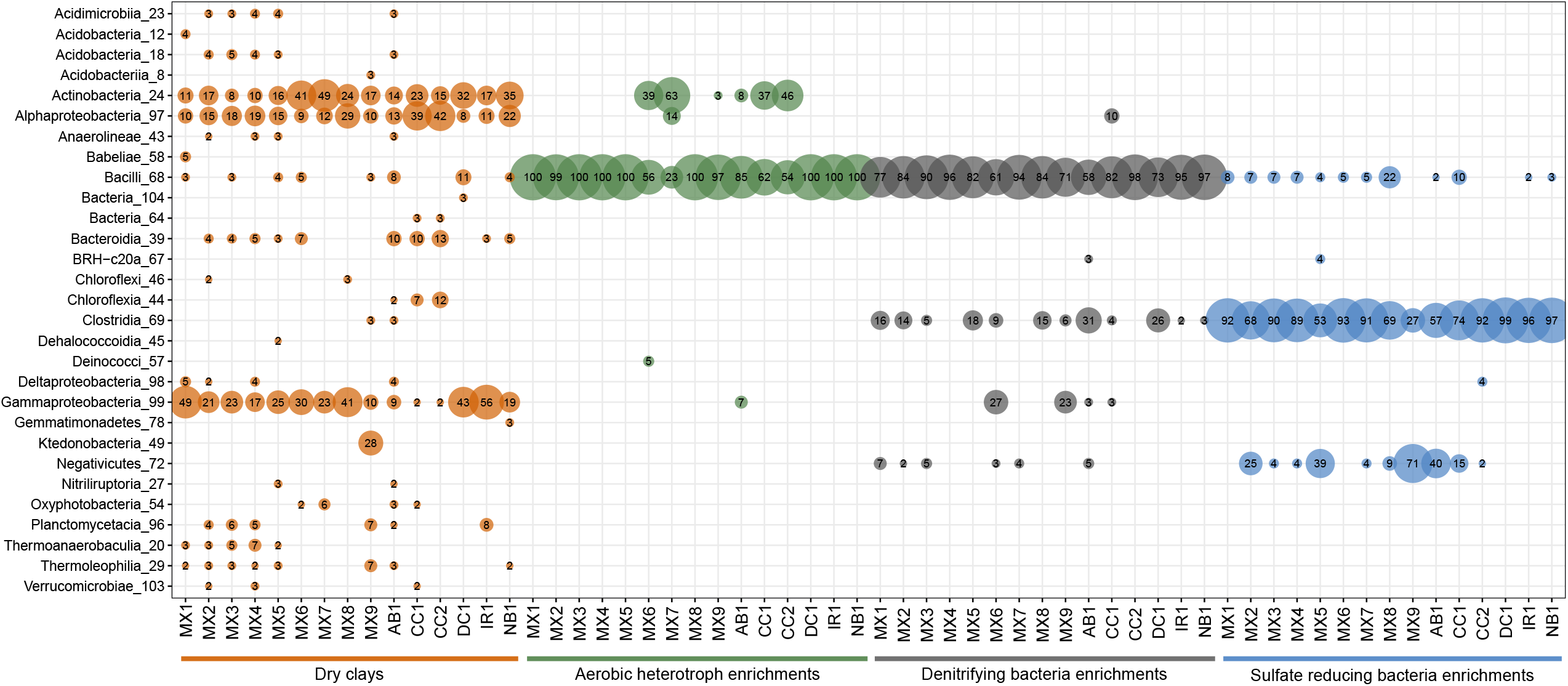
Bubble plot of ASV taxonomic affiliations collapsed to the class level for merged replicates of enrichment culture and as-received clay samples. All orders at or above 2% relative abundance for a merged replicate sample are shown.

The microbial profiles generated from enrichment cultures of denitrifying bacteria, SRB, and aerobic heterotrophs included overlapping ASVs associated with bacteria belonging to the genera *Bacillus, Clostridium*, and *Paenibacillus* (Figures S5, S6, and S7). Similarities between the ASVs in the 16S rRNA gene profiles of the denitrifying bacteria and SRB enrichments were likely due to the same culturable microorganisms from the clays responding positively to similar cultivation conditions that supported anaerobic heterotrophy. Compared to ASVs associated with *Bacillus* and *Clostridium*, ASVs associated with *Pseudomonas* were detected less frequently in denitrifying enrichment cultures (Figure S6). *Pseudomonas* spp. are commonly reported in enrichment cultures of natural clay deposits or saturated and highly compacted clays (16,26,38– 41), and have sometimes been reported in uncompacted as-received clay (18). Several ASVs associated with the genera *Paracoccus* and *Thiobacillus*, potentially capable of denitrification and sulfur oxidation respectively (8,10,42), were detected in as-received clays but not in enrichment cultures, although they have also previously been detected in clay enrichment cultures (13,26).

Enrichment cultures of sulfate-reducing bacteria promoted growth of bacteria from genera such as *Bacillus, Desulfosporosinus, Desulfitobacterium, Clostridium, Anaerosolibacter*, and *Sedimentibacter* (Figure S7). *Desulfosporosinus*, a common genus of sulfate-reducing bacteria (42), was cultivated in SRB enrichments from every bentonite sample, but was rarely detected at a relative abundance greater than 2% in the initial clay samples (Figure 4). In previous research, *Desulfosporosinus* spp. were also frequently detected in microcosms and enrichment cultures from multiple types of bentonite clay (13,18,39). Although many studies suggest that SRB make up the largest group within the microbial communities of clays (11,12,43,44), and SRB were dominant in SRB enrichments here, microbial community profiles of initial clay DNA extracts were not dominated by SRB (Figure 6). *Pseudomonas* and *Desulfosporosinus* ASVs detected in our enrichment cultures were identical to several of those from compacted bentonite exposed to natural groundwater in borehole modules (38), which indicates that the results presented here have direct “real world” implications for a proposed deep geological repository.

This cultivation data implies that similar taxa to those cultured here may respond and grow in a nuclear repository environment if the appropriate engineered barrier component design conditions are not established. This also indicates that nuclear waste management organizations can effectively choose any mined bentonite clay material that achieves a sufficient swelling pressure when saturated, regardless of origin or initial molecular microbial profiles, because they all possess core taxa that respond similarly and opportunistically when conditions become suitable.

### Core microbial community

Assessments of microbial profiles obtained from enrichment cultures and as-received clay DNA extracts can provid conflicting perspectives of the relative abundance and viability of clay microorganisms. Although sequencing of enrichment culture DNA revealed clay microorganisms that were viable, this approach did not measure the absolute abundance of microorganisms within clay samples. For direct comparisons of microorganisms detected from enrichment cultures and as-received clay DNA extracts, each ASV was categorized as being present only in enrichment cultures, only in clay, or present in both enrichment cultures and as-received clays. Of those ASVs, 81.9% (1604 ASVs) were only detected in clays, 16.8% (330 ASVs) only in enrichment cultures, and 1.3% (25 ASVs) were detected in both, highlighting that very few taxa were both cultivable and present in abundances sufficient for detection in direct DNA extracts from as-received clays (Figure 7A). As resported by studies exploring soil and the human gut, cultivation preferentially recovers microorganisms of low relative abundance, often from the so-called “rare biosphere” (45,46).

**Figure 7.**
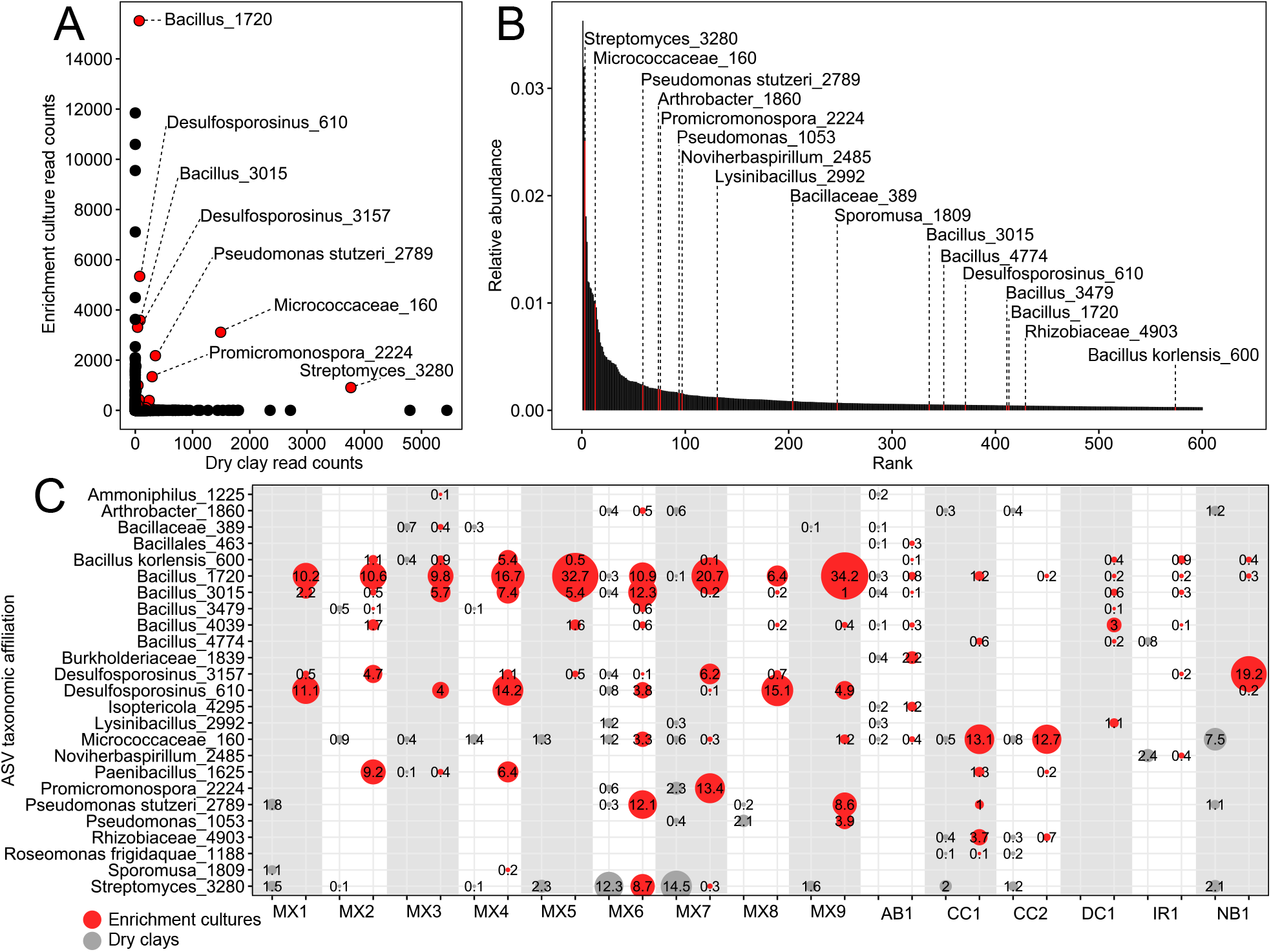
Core microbial community detected in bentonite clays after normalization to 10,000 reads. (A) Scatter plot of 16S rRNA gene sequence read counts from clays and enrichment cultures. Core ASVs, detected in both clays and enrichment cultures, are indicated with a red point. (B) Rank abundance curve of ASVs detected in clays, highlighting ASVs additionally detected in enrichment cultures in red. (C) Bubble plot of core clay ASVs present in clays and enrichment cultures. The bubbles represent ASVs present at ≥0.1% relative abundance in samples and the numbers contained within specify the percentages. For ASV labels, we report the lowest taxonomic ranks that have confidence values above the default 0.7 threshold. For each sample, sequences obtained from duplicates of all three enrichment cultures or from duplicate clay samples were merged into enrichment culture or as-received clay categories, respectively.

Of the 600 most abundant ASVs directly detected within all dry clays, most were only detected in DNA extracts from as-received clay (Figure 7B), indicating that they represent free DNA, slow-growing taxa, or microorganisms that did not respond to the culturing conditions used in this study. The 25 most abundant ASVs detected in both clay and enrichment cultures classified confidently to the following taxonomic ranks *Streptomyces, Micrococcaceae, Promicromonospora, Bacillus, Rhizobiaceae, Pseudomonas, Burkholderiaceae, Desulfosporosinus, Noviherbaspirillum*, and *Isoptericola* (Figure 7C). Detected in 10 of the 15 clay samples, *Streptomyces* was the most abundant ASV from all as-received clay DNA extracts combined (Figure 7B), although five samples (AB1, IR1, DC1, MX3, and MX8) did not contain detectable *Streptomyces* with relative abundances greater than 0.1% (Figure 7C). *Streptomyces* was also detected in enrichment cultures of two samples (MX6 and MX7; Figure 7C) indicating the viability of these microorganisms in clay. Previous studies of microorganisms within compacted bentonite corroborate the presence (38,40) and viability (40) of *Streptomyces*. Species of *Micrococcaceae* and *Bacillus* were previously detected in compacted bentonite (38), natural bentonite formations (47), and clay enrichment cultures (18), and were also detected here in as-received dry bentonite samples and their corresponding enrichment cultures (Figure 7). The large proportion of *Micrococcaceae* and *Bacillus* in many enrichment cultures, compared to their corresponding dry clay samples, confirms that representatives were viable and cultureable bacteria, prevalent across samples, making them core members of diverse bentonite clays.

## Conclusion

Microbial 16S rRNA gene profiles of as-received clay samples revealed distinct microbial community profiles but dominant ASVs did not reflect the viable bacteria enumerated with cultivation-dependent approaches. Enrichment cultures routinely selected for the same taxa regardless of clay starting material. The ASVs that were detected in both as-received bentonite clay DNA and enrichment culture DNA profiles were primarily associated with desiccation-resistant taxa, potentially surviving in the dry clay through formation of spores or by preservation within the bentonite clay, including those affiliated with *Streptomyces, Micrococcaceae, Bacillus*, and *Desulfosporosinus*. Detection of ASVs in both enrichment culture and as-received clay implies that the associated bacteria were viable in clay, and therefore are key members of the overlapping core culturable community subset and core microbiome of bentonite clays. Identifying the common microbial “targets” that can grow when conditions are suitable is important because this informs ongoing bentonite experiments that simulate saturated DGR barrier conditions. For example, detection of the same target microorganisms in all clay samples allows us to conclude that experimental results using one bentonite clay type may be extrapolated more generally to others as well. Future research should explore whether abundant taxa detected within 16S rRNA gene profiles that were not recovered in cultures are viable but uncultureable or instead reflect relic DNA that is adsorbed to the charged clay matrix. Given that microbiology is core to building a safety case for repository design, this study will be critical for nuclear waste management organizations globally as research continues investigating the microbiology of engineered barrier components for a deep geological repository.

## Materials and Methods

### Clay sample selection

In order to obtain a diverse subset of bentonite clay, our samples were selected from four different countries, with five different manufacturers, and included bentonite clays dominated by either sodium or calcium exchangeable cations (Table 1). In general, following excavation of bentonite from a deposit, manufacturers crush and dry the crude ore to around 20 – 25% moisture content by mass (48). Bentonite can either be air dried in stockpiles or dehydrated in a rotary drier prior to storage. Screening and mixing of the stockpiles is also commonly conducted to achieve a specific grade (i.e., particle size distribution and chemistry) of material. After receiving the samples, all coarse as-received clay samples (i.e., MX7) were ground to a fine grain size using a DNA-free glass mortar and pestle. Two bentonite samples were provided with different initial granularities (CC1 and CC2, and MX6 and MX7) but, even though technically replicates, were treated as separate samples in this study. Wyoming MX-80 samples with different production dates and lot numbers were used to investigate possible influences of batch characteristics on microbial community profiles (Table 1).

### Moisture content and water activity

Water potential was measured using a WP4 Dew Point Potentiometer (Meter Group, USA) with 2-5 g of as-received clay, following the manufacturer instructions for “fast mode” analysis at 25.0°C. Water activity was calculated according to manufacturer instructions, using the potentiometer output of pressure (kPa) and temperature (°C). Moisture content was calculated by measuring the loss of water after heating clay at 110°C for 24 hours and weighing samples before and after drying using the following formula: (g_wet_ – g_dry_) / g_wet_.

### Cultivation of bentonite clay bacterial communities

A dilution series was prepared in sterile phosphate-buffered saline solution (PBS; 0.01 M NaCl buffered to pH 7.6 with 9 mM Na_2_HPO_4_ and 1 mM NaH_2_PO_4_) with clay dilutions of 10^−1^-10^−3^. The 10^−1^ dilution was prepared in a 50-mL conical tube by slowly adding 2 g of clay to 18 mL of PBS while vortexing continuously, followed immediately by continuous gentle agitation for 30 minutes at room temperature. The agitation time was necessary to allow the clay to suspend and swell evenly, which has been shown to result in greater homogeneity of the clay-PBS solution and a higher efficiency of cell removal from clay interfaces (27). Remaining dilutions were prepared by transferring 1 mL of the previous dilution into 9 mL of PBS. Aliquots from all dilutions were dispensed into most-probable number (MPN) test tubes (1 mL inoculum into 9 mL of medium) and onto R2A agar spread plates (100 μL inoculum) as described previously (24).

For enrichment and enumeration of SRB, MPN test tubes were prepared with 9 mL of sterile sulfate-reducing medium (HiMedia Laboratories, M803). For denitrifiers, MPN tubes with liquid R2A medium (HiMedia Laboratories, M1687) were amended with 12 mM sodium nitrate and included an inverted Durham tube. For SRB and denitrifiers, each sample was analyzed using a five tube MPN method, with all test tubes placed into a stainless steel vacuum chamber (BVV, USA) containing a GasPak EZ Anaerobe Container System Sachet (BD, USA) and an anaerobic indicator strip (BD). Culture chambers were evacuated and flushed with N_2_ before incubation for 28 days at 30°C. After incubation, positive MPN tubes were identified by a black precipitate for SRB or by a gas bubble in the inverted Durham tube for denitrifying bacteria activity. The MPN per gram dry weight (gdw) was calculated according to the moisture content measured for each sample. Mean MPN/gdw and standard deviation values were calculated based on triplicate MPN assays.

Aerobic and anaerobic heterotrophs were cultured on R2A medium with 1.5% agar (29). Plates were incubated at 30°C under oxic conditions for 5-7 days or under anoxic conditions for 28 days. Colony forming units (CFU)/gdw were calculated using the sample moisture contents, and standard deviation values were calculated based on three replications of each plate count analysis. The lower limit of quantification for heterotrophic plate counts was 2500 CFU/g because plates with fewer than 25 colonies may exaggerate low cell counts (49).

### Genomic DNA extraction from clays and enrichment cultures

Genomic DNA was extracted from 2 g powdered clay samples using the PowerMax Soil DNA Isolation Kit (Qiagen, Germany) and modifications to the manufacturer’s instructions previously validated (22); after addition of lysis solution, clay in PowerBead tubes was gently vortexted for 20 minutes to allow clay to fully suspend and swell, then PowerBead tubes were incubated at 65 °C for 30 minutes, immediately followed by bead beating at 30 Hz for 10 minutes using a mixer mill MM 400 (Retsch). Kit controls were included for each batch of DNA extractions.

For extraction of enrichment culture genomic DNA, colonies on replicate agar plates for one sample from the same dilution were slurried by adding 1 mL of sterile DNA-free water and gently sweeping over the plate surface with a sterile disposable cell spreader. The slurry was then transferred to a DNA-free microcentrifuge tube. The contents of positive MPN tubes were mixed, then 2 mL of the culture was transferred to a DNA-free microcentrifuge tube. Cells were pelleted by centrifuging all microcentrifuge tubes for 2 minutes at 10,000 x g. Genomic DNA was recovered from cell pellets following the protocol for the DNeasy Ultraclean Microbial Kit (Qiagen) using a bead beater (FastPrep-24 Instrument MP Biomedicals, USA) at 5.5 m/s for 45 seconds. The DNA from replicate MPN tubes of the same dilutions were pooled before amplification.

### Quantitative polymerase chain reaction (qPCR)

All qPCR mixtures were prepared in a PCR hood (AirClean Systems, Canada) that was cleaned with 70% ethanol and treated with UV light for 15 minutes. The qPCR standard curve template was generated from the V3-V5 16S rRNA gene fragment of *Thermus thermophilus* that was previously cloned into vector pUC57-Kan. The template was amplified through PCR with primers M13F (5’-TGTAAAACGACGGCCAGT-3’) and M13R (5’-CAGGAAACAGCTATGAC-3’) that flanked the 719 bp insert. The PCR product was separated on a 1% agarose gel and purified using the Wizard SV Gel and PCR Clean-Up System (Promega, USA).

Genomic DNA extracts from clay samples were quantified using universal 16S rRNA gene primers 341F and 518R (50). All qPCR amplifications were performed in duplicate and each contained 15 µL total volume of 1 × SsoAdvanced Universal SYBR Green Supermix (Bio-Rad, USA), 0.3 µM of each primer, 7.5 µg bovine serum albumin (BSA), and 4 µL of template DNA. The qPCR amplification was performed with a CFX96 Real-Time PCR detection system (Bio-Rad) beginning with 98°C for 3 minutes followed by 40 cycles of 98°C and 55°C at 15 s and 30 s intervals, respectively. Initial 16S rRNA gene copy numbers were calculated for kit controls and clay samples from the linear regression equation produced from the standard curve with a 0.98 coefficient of determination (*R*^2^). Average starting quantities of up to 1.5×10^3^ copies/mL detected in kit controls were subtracted from the calculated starting quantities for respective samples. Final 16S rRNA gene copy numbers were corrected to per gram dry clay values using the moisture content measurements of each sample.

### Amplification of 16S rRNA genes and high-throughput sequencing

All PCR mixtures were prepared in a PCR hood (AirClean Systems, Canada) that was cleaned with 70% ethanol and treated with UV light for 15 minutes. Each 25 μL PCR mixture contained 1× ThermoPol Buffer, 0.2 μM forward primer, 0.2 μM reverse primer, 200 μM dNTPs, 15 μg BSA, 0.625 units of hot start *Taq* DNA polymerase (New England Biolabs, USA), and up to 10 ng of template DNA. The V4-V5 region of the 16S rRNA gene was amplified in triplicate using primers 515F-Y (51) and 926R (52), modified to contain unique 6 base indexes in addition to Illumina flow cell binding and sequencing sites (53). Reaction conditions were an initial denaturation at 95°C for 3 min, followed by 40 cycles of denaturation at 95°C for 30 sec, annealing at 50°C for 30 sec, and extension at 68°C for 1 min, with a final extension at 68°C for 7 min. Negative controls containing no template DNA (NTCs) were included in the 96 well PCR plates to test for cross contamination, and in tubes outside of the PCR plate to test for master mix contamination.

Based on agarose gel quantification, samples were normalized by pooling equimolar quantities into a single tube. The NTCs and DNA extraction kit controls were added to the pool with 5 µl each even if no visible band was detected on the agarose gel. The pooled amplicons were gel purified using the Wizard SV Gel and PCR Clean-Up System (Promega, USA) and the library was denatured and diluted following manufacturer’s guidelines (Illumina document no. 15039740 v10). The 8 pM library containing 15% PhiX control v3 (Illumina Canada, Canada) was sequenced on a MiSeq instrument (Illumina Inc, USA) using a 2 × 250 cycle MiSeq Reagent Kit v2 (Illumina Canada). Samples were sequenced in two MiSeq runs and reads were merged in the post sequence analysis.

### Post sequencing analysis

Sequence reads were demultiplexed using MiSeq Reporter software version 2.5.0.5 (Illumina Inc) and analyzed using Quantitative Insights Into Microbial Ecology 2 (QIIME2; version 2019.10.0) (54) that denoises sequences with DADA2 (release 1.16) (55). Samples were rarefied to 2,350 sequences for generating ordinations and collapsed ASV tables were generated using QIIME2 plugins. These analyses were managed by Automation, Extension, and Integration Of Microbial Ecology version 3 (AXIOME3; <u>github.com/neufeld/AXIOME3</u>) (56) and taxonomy was assigned to ASVs using SILVA 132 release (57–59). Decontam (release 3.11) was used to identify contaminant ASVs using the prevalence method and an assigned score statistic threshold value of 0.5 (58). These ASVs were verified and removed manually from the sample ASV table and summarized in Figure S4. Controls used for the Decontam analysis included a swab from sterile R2A agar plate, kit controls for DNA extraction (3 controls), PCR no-template controls (NTCs; 8 controls), and positive controls for sequencing (3 controls). Next, the microbial community profiles detected from each dry clay sample was compared to the profiles detected after enrichment culturing. For this comparison, the ASVs detected in each sample’s three different enrichment cultures were added together as one enrichment culture category. All ASV read counts were normalized to 10,000 reads for each sample. From all of these normalized reads, ASVs representing ≥ 0.1% relative abundance in the dry clay microbial profile of ≥1 sample and in the enrichment culture microbial profile of ≥1 sample were labelled as core ASVs.

### Phospholipid fatty acid analysis

Phospholipid fatty acid (PLFA) analysis was carried out by Microbial Insights (Knoxville, TN). Lipids were recovered using a modified Bligh and Dyer method (61). Estimates of prokaryotic cells per gram based on PLFA were calculated with the conversion of 20,000 cells/pmol of PLFA (62). To allow cross comparison with other prokaryotic enumerations, eukaryote-associated PLFA quantities were removed and only prokaryotic PLFA quantities were reported for estimates of cell abundance. Each category of PLFA is generally associated with specific groups of microorganisms, except for Nsats which are found in all organisms. The PLFA analyses were performed in singlicate, but duplicates were analyzed for samples AB1 and IR1.

All DNA sequences were deposited in the European Nucleotide Archive with study accession number PRJEB39383.

## Acknowledgements

We thank Zinedine Cobourne for assistance with cultivation and Daniel Min for help with sequence analysis. This research was supported by an Ontario Research Fund: Research Excellence (ORF-RE) grant and a Collaborative Research and Development (CRD) grant from the Natural Sciences and Engineering Research Council of Canada (NSERC).

