## Supplemental material for "Fifteen shades of clay: distinct microbial community profiles obtained from bentonite samples by cultivation and direct nucleic acid extraction"

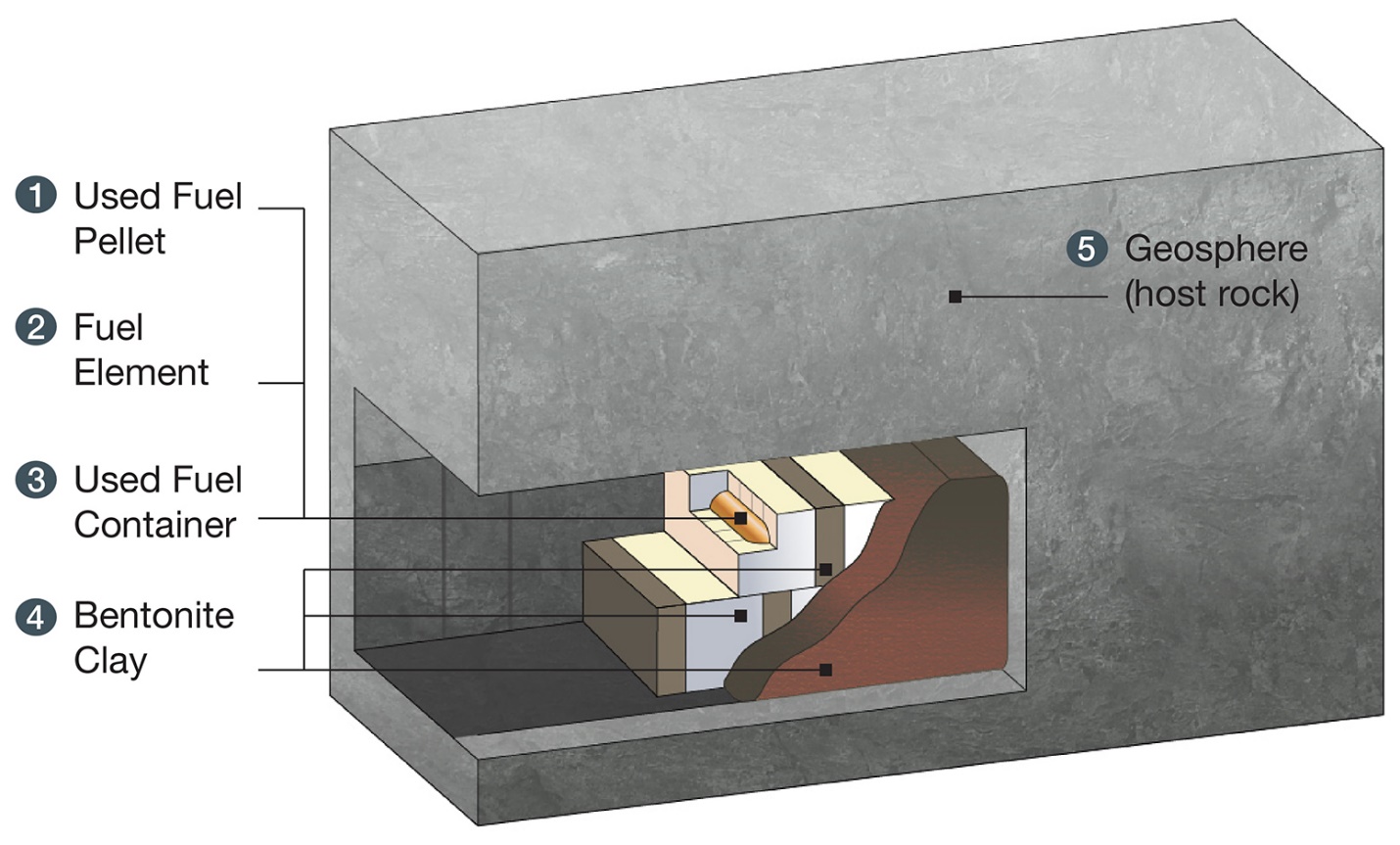

Supplemental figure 1. A schematic representation of an underground emplacement room proposed for the Canadian DGR for used nuclear fuel. The used fuel and fuel element are sealed in a copper-coated carbon steel used fuel container and placed in a highly compacted bentonite box approximately 500 m below ground. Granular bentonite is used to fill the gaps left between the rock wall and the compacted bentonite.

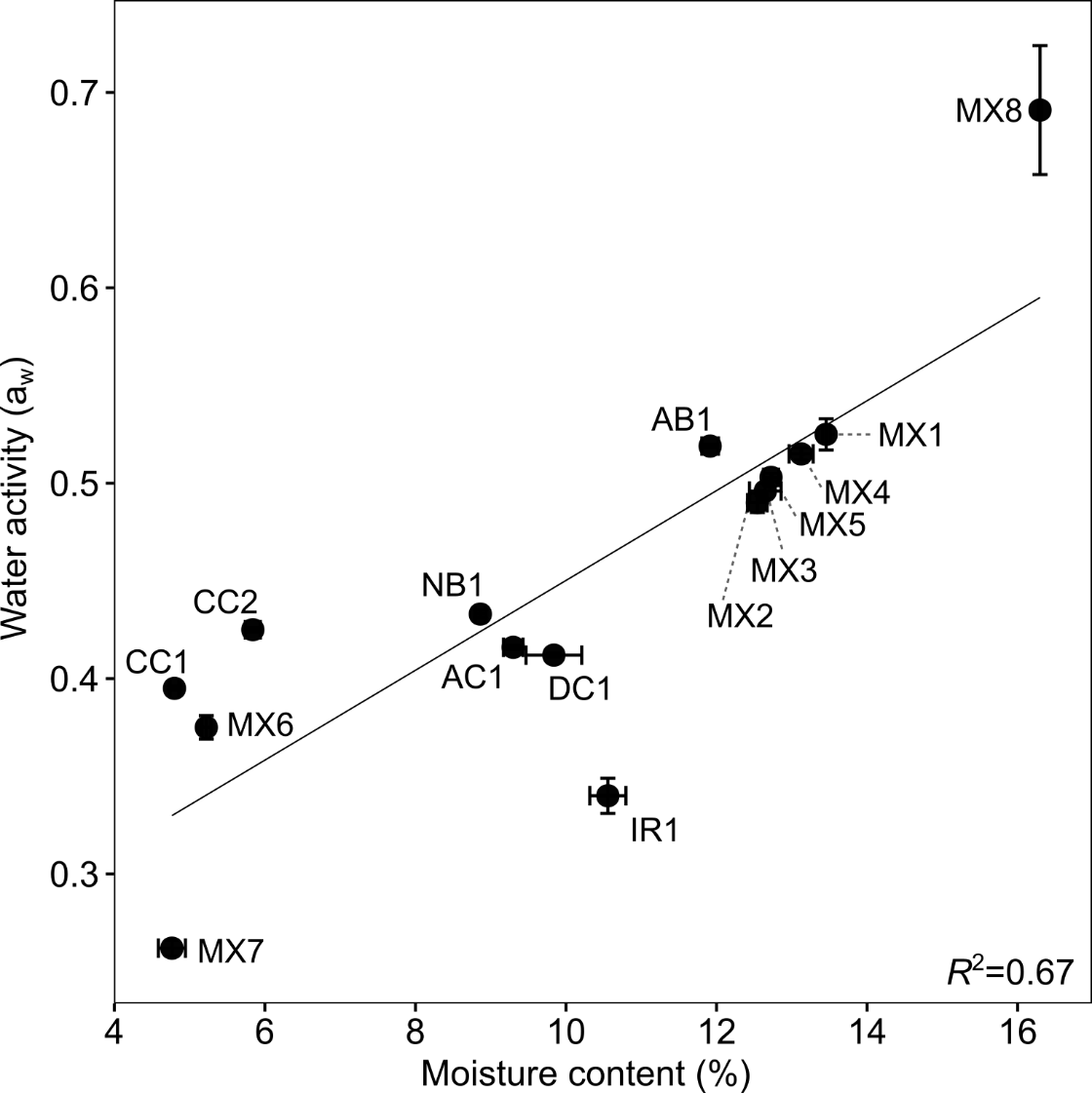

Supplemental figure 2. Moisture content and water activity plot for all as-received clay samples. Error bars represent standard deviation of triplicate analyses and the solid line is the linear regression line for samples. Water activity calculated using a water potentiometer and moisture content values calculated after evaporation of moisture in an oven.

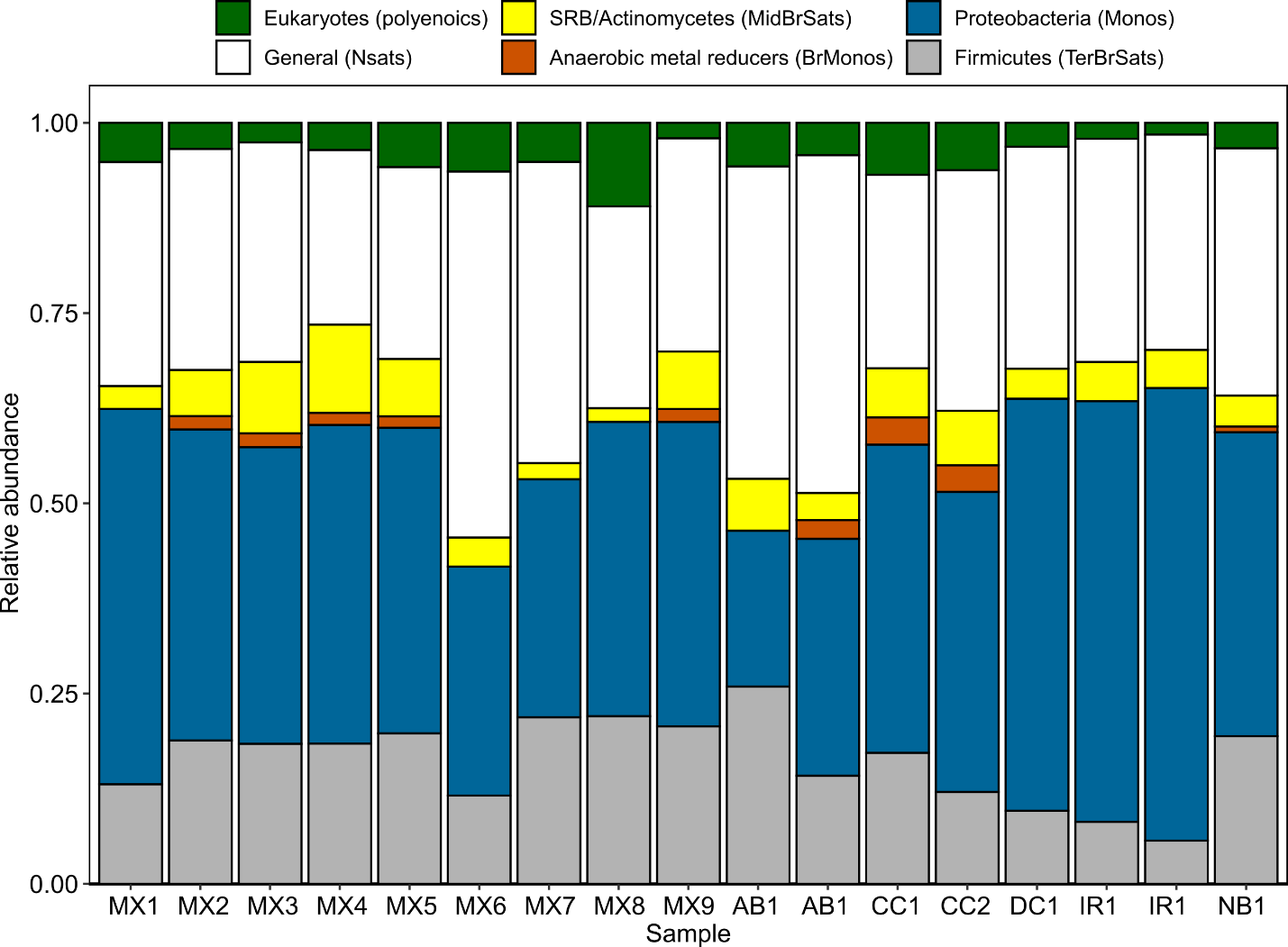

Supplemental figure 3. Phospholipid fatty acid (PLFA) based community structure analysis across diverse clay samples. Structural groups are separated into different categories based on monoenoic (Monos), terminally branched saturated (TerBrSats), branched monoenoic (BrMonos), mid-chain branched saturated (MidBrStats), normal saturated (Nsats), and polyenoic polyenoic PLFA profiles detected in each sample.

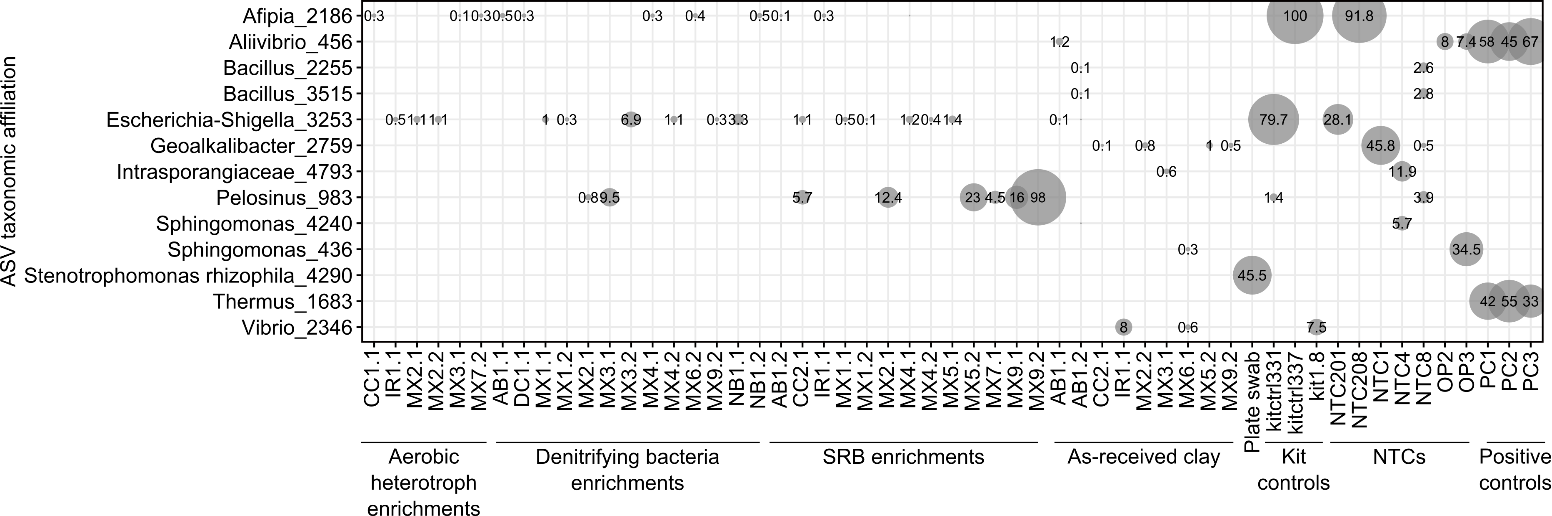

Supplemental figure 4. Contaminant ASVs identified in enrichment culture and as-received clay samples. Only contaminants ≥ 0.1% relative abundance in samples are pictured. Sample names include the sample ID, the source of DNA (A, aerobic heterotroph enrichments; D, denitrifying bacteria enrichments; S, SRB enrichments; X, dry clay) and the replicate number (1 or 2). For ASV labels, we report the lowest taxonomic ranks that have confidence values above the default 0.7 threshold. The presented ASVs were removed from the ASV table before further analysis was performed. Negative controls include a “plate swab” from the surface of uninoculated R2A agar, kit controls used for each batch of DNA extractions, and no-template controls (NTCs).

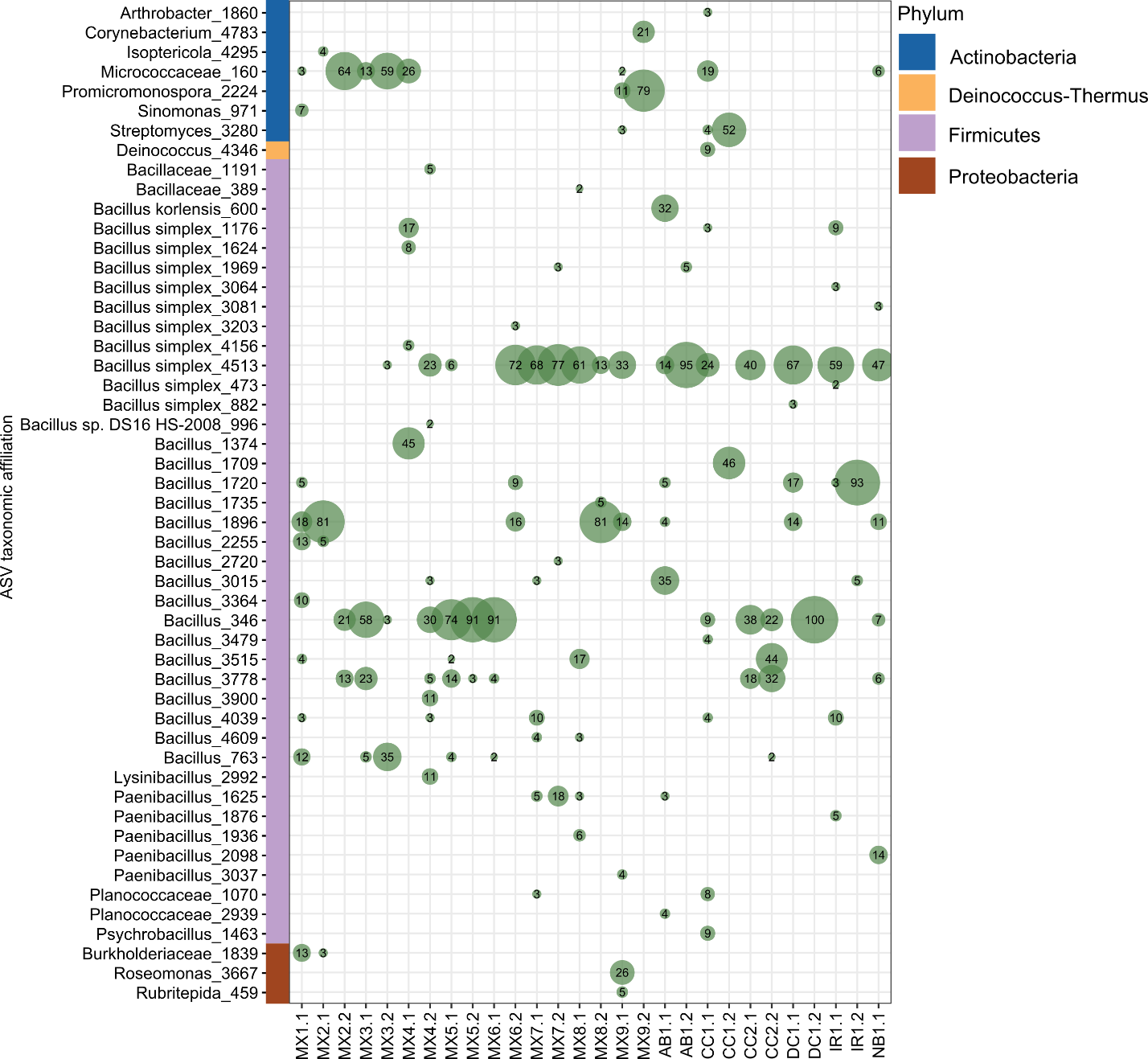

Supplemental figure 5. Relative abundances of ASVs affiliated with enrichment cultures for aerobic heterotrophs from clay samples. Duplicate sample community profiles represented at the ASV level with a relative abundance ≥2%. For ASV labels, we report the lowest taxonomic ranks that have confidence values above the default 0.7 threshold. Sample replicate number is indicated at the end of each sample name.

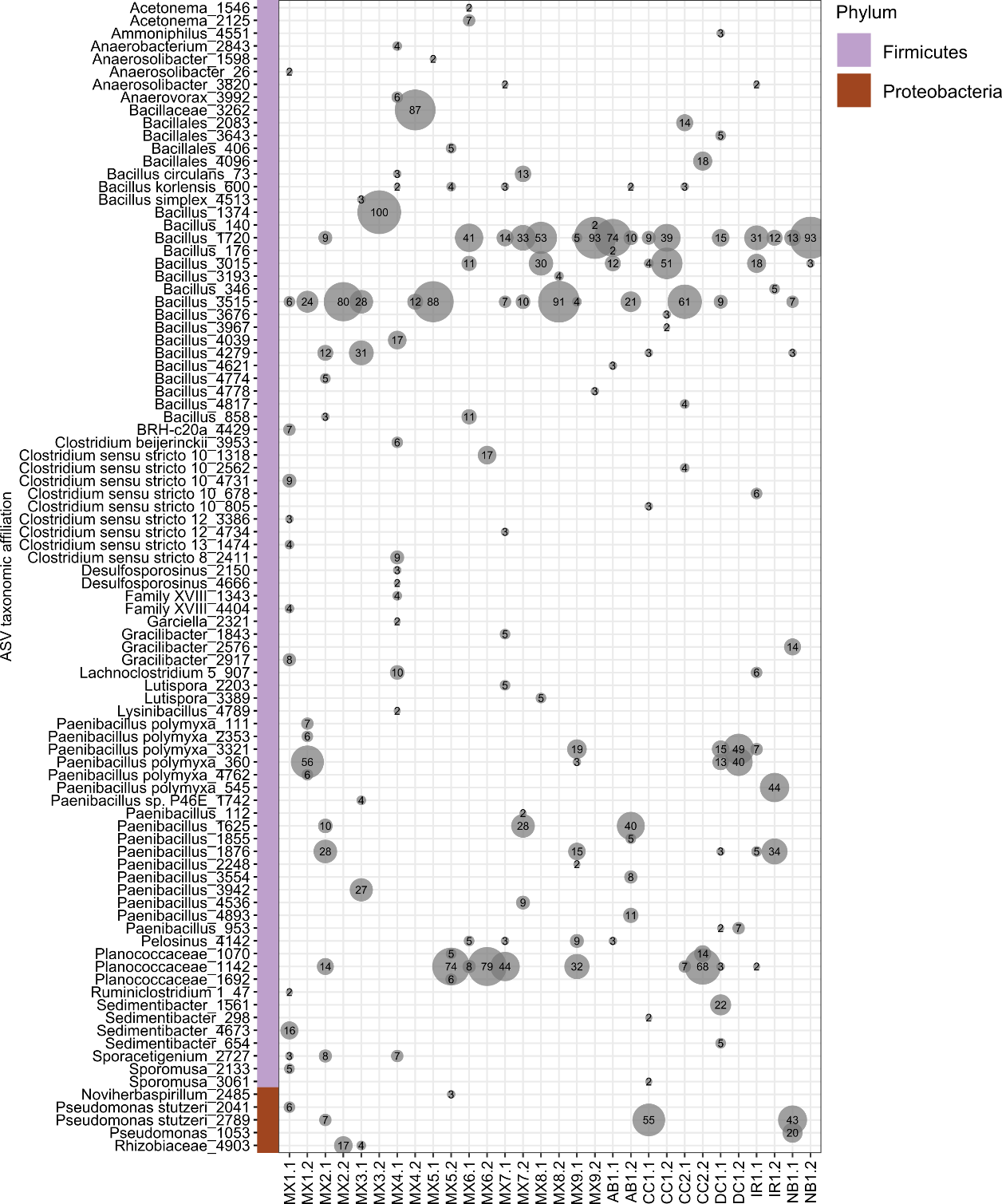

Supplemental figure 6. Relative abundances of ASVs affiliated with enrichment cultures for denitrifying bacteria from clay samples. Duplicate sample community profiles represented at the ASV level with a relative abundance ≥2%. For ASV labels, we report the lowest taxonomic ranks that have confidence values above the default 0.7 threshold. Sample replicate number is indicated at the end of each sample name.

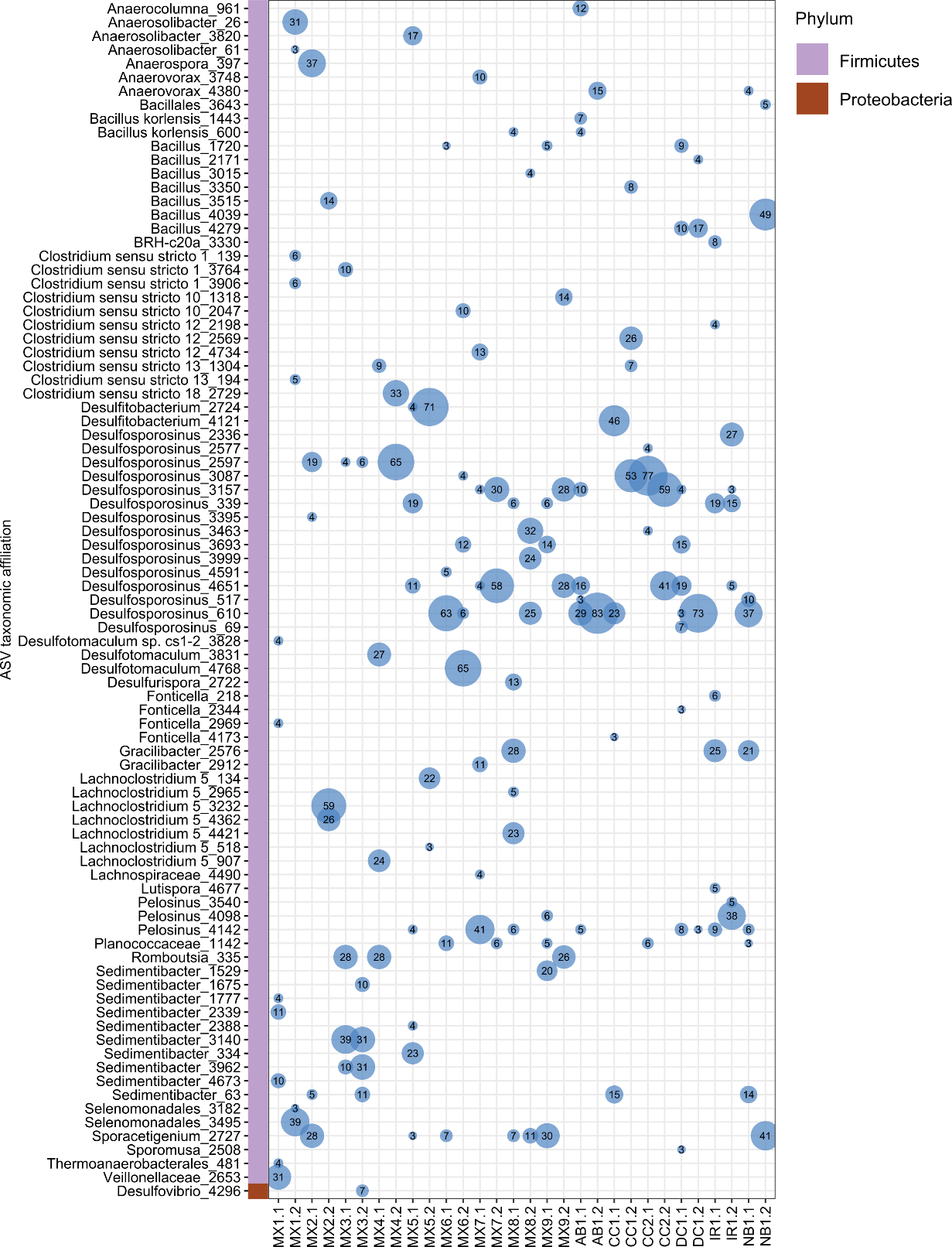

Supplemental figure 7. Relative abundances of ASVs affiliated with enrichment cultures for sulfate-reducing bacteria from clay samples. Duplicate sample community profiles represented at the ASV level with a relative abundance ≥3%. For ASV labels, we report the lowest taxonomic ranks that have confidence values above the default 0.7 threshold. Sample replicate number is indicated at the end of each sample name.
